# Patch-Clamp Proteomics of Single Neuronal Somas in Tissue Using Electrophysiology and Subcellular Capillary Electrophoresis Mass Spectrometry

**DOI:** 10.1101/2021.09.02.458040

**Authors:** Sam B. Choi, Abigail M. Polter, Peter Nemes

## Abstract

Understanding of the relationship between cellular function and molecular composition holds a key to next-generational therapeutics but requires measurement of all types of molecules in cells. Developments in sequencing enabled semi-routine measurement of single-cell genomes and transcriptomes, but analytical tools are scarce for detecting diverse proteins in tissue-embedded cells. To bridge this gap for neuroscience research, we report the integration of patch-clamp electrophysiology with subcellular shot-gun proteomics by high-resolution mass spectrometry (HRMS). Recording of electrical activity permitted identification of dopaminergic neurons in the substantia nigra pars compacta. Ca. 20–50% of the neuronal soma content, containing an estimated 100 pg of total protein, was aspirated into the patch pipette filled with ammonium bicarbonate. About ~1 pg of somal protein, or ~0.25% of the total cellular proteome, was analyzed on a custom-built capillary electrophoresis (CE) high-resolution mass spectrometer (HRMS). A series of experiments were conducted to systematically enhance detection sensitivity through refinements in sample processing and detection, allowing us to quantify ~275 different proteins from somal aspirate-equivalent protein digests. From single neurons, patch-clamp proteomics of the soma quantified 91, 80, and 95 different proteins from 3 different dopaminergic neurons, or 157 proteins in total. Quantification revealed detectable proteomic differences between the somal protein samples. Analysis of canonical knowledge predicted rich interaction networks between the proteins. The integration of patch-clamp electrophysiology with subcellular CE-HRMS proteomics expands the analytical toolbox of neuroscience.

## INTRODUCTION

Understanding the relationship between cellular function and molecular composition holds a key to better diagnoses and therapeutics but requires major advances in single-cell analyses. In the study of the brain, direct measurement of cellular electrical activity supports physiology, charting of cellular phenotypes, and eavesdropping on inter-cellular communication.^1–3^ Concurrent detection of all the molecules produced by neurons, ranging from transcripts and proteins to peptides and metabolites, can help us gauge molecular states during states of homeostasis or imbalance. Because proteins are sensitive to both intrinsic and extrinsic events, the single-cell proteome promises to be an effective indicator of the overall molecular state of the cell. Patchclamp electrophysiology was combined with single-cell and subcellular sequencing of transcripts^1,4^ and high-resolution mass spectrometry (HRMS) of peptides and metabolites^5–9^. Information on the proteome of physiologically characterized neurons, however, is lacking, even today. There is a high and still unmet need for designing technologies capable of characterizing broad types of proteins in electrophysiologically characterized cells, ideally in their native tissue environment using HRMS.

Single-cell HRMS enabled molecular characterization of the cell with high sensitivity and molecular specificity. The current state of this field was the focus of several reviews.^10–17^ Recent technologies, such as nanoPOTS^18,19^, oil-air-droplet (OAD) chip^20^, and integrated proteome analysis device (iPAD)^21^, advanced microdroplet processing to analyze limited amounts of protein digests from single cells. With these approaches, together with specialized multiplexing strategies, such as SCoPE^22^, it is now possible to analyze hundreds of different proteins in sorted cells or chemically fixed tissues. Biological and translational studies requiring information on the physiological state of electrically active cells or cell compartments, such as neurons, would greatly benefit from concurrent analysis of the proteomic state of the system.

Capillary electrophoresis (CE) equips single-cell MS with scalability (reviewed in Ref. ^23^). Analytical volumes in CE are compatible with limited sample amounts from single cells. The technology enhances HRMS detection sensitivity with exquisite separation power and various methods for on-column enrichment.^24,25^ An international study recently found CE-HRMS robust across laboratories.^26^ We^27–29^ and others^30,31^ custom-built microanalytical CE platforms and built ultrasensitive CE electrospray ionization (ESI) interfaces (reviewed in Ref. ^23^) to quantify transcripts, proteins, peptides, and metabolites in single cells ex vivo^29,32^, *in situ*^33,34^, or *in vivo*^33^.These studies yielded new tools for cell, developmental, and neurobiology and led to the discovery of molecules capable of altering normal tissue developmental trajectories.^28^

These performance metrics by CE-MS proteomics are attractive for patch-clamp electrophysiology. Recently, we custom-built CE-MS platforms with a capability for ~250 zmol lower limit of detection for model peptides^35,36^. These instruments supported the identification of ~200–800 proteins in protein digests diluted to estimate to the total composition of ~5–10 neurons.^35,37,38^ A data-dependent acquisition ladder enhanced these limitations, identifying 428 proteins from single-cell equivalent protein amounts from cultured neurons.^38^ Based on these technical performance metrics, we recently explored a technical capability to combine electrophysiology with single-neuron CE-MS proteomics.^39–41^

Here, we present a systematic study that enabled the combination of patch-clamp electrophysiology with subcellular CE-ESI-MS to quantify proteins in identified neurons in brain tissues. Inspired by the success of patch-clamp single-cell transcriptomics^1,4^, peptidomics^5^, and metabolomics^6–8^, in 2019, we reported the use of a patch pipette as a microprobe to bridge electrophysiology with microsampling for CE-MS proteomics.^39–41^ Here, we report additional refinements, testings, and validations of the method to enhance performance metrics. The approach was developed using dopaminergic (DA) neurons in sections of the substantia nigra pars compacta (mouse), because these cells were identifiable based on size, location, and electrical activity. After patch-clamping the neurons under an upright microscope, cellular phenotype was identified based on the detection of slow pacemaker activity. Approximately 20–50% of the neuronal soma, containing ~100–250 pg of total protein amount, was aspirated into the patch pipette. The extracted proteins were processed to 3 μL of tryptic digest following a workflow that we developed to systematically enhance protein detection and quantification using MS. About 1 pg, or ~0.25% of the total protein extracted from the neuron, was analyzed on an ultrasensitive CE-ESI platform using a quadrupole-orbitrap hybrid mass spectrometer. Quantification of 157 different proteins between the somal aspirates revealed detectable variability in their proteomes. Integration of patch-clamp electrophysiology with subcellular CE-HRMS proteomics expands the analytical toolbox of neuroscience.

## EXPERIMENTAL SECTION

### Materials

All materials were purchased at reagent grade or higher. Samples for CE-MS were prepared in LC-MS quality solvents. Further details are available in the electronic **Supplementary Information (SI)** document.

### Animals and Preparation of Brain Sections

All procedures were carried out in accordance with the guidelines of the National Institutes of Health for humane animal care and use under approval by the George Washington University Institutional Animal Care and Use Committee (Approval no. A378). Acute brain tissue sections were vibratome-sectioned from C57Bl6/J mice following standard protocols. Specifics are in the **SI** document.

### Whole-Neuron Electrophysiology and Sample Collection

Midbrain slices were continuously perfused with artificial cerebrospinal fluid. Electrical recordings were conducted in an electrophysiology system equipped with an upright microscope (FN1, Nikon USA, Melville, NY). Patch pipettes for recording (2–4 MΩ) and protein extraction were backfilled with ~20 μL 50 mM ammonium bicarbonate (AmBic), unless otherwise specified. DA neurons were identified based on slow pacemaker activity, morphology, and location in the tissue. Ca. 20–50% volume of the soma volume was aspirated into the patch pipette. The samples were processed for shot-gun proteomics. Multiplexing quantification used the TMT-128 and TMT-131 channels from a 6-plexing tandem mass tag (TMT) kit (Thermo Fisher, Waltham, MA). See details in the **SI** document.

### Proteomics by CE-ESI-HRMS

The neuron samples were analyzed on a microanalytical CE-ESI-HRMS system that we custom-built and validated for ultrahigh sensitivity following our recent studies.^35,37,38^ Peptides from the somal protein digests were electrophoresed in 25% acetonitrile (ACN) with 1 M formic acid (background electrolyte), then ionized in a coaxial low-flow sheath-flow CE-ESI interface. This CE-ESI platform was constructed based on an independent design^42,43^, which we operated in the cone-jet spraying regime^44^ to maximize detection sensitivity.^37,38^ Peptide ions were detected on a hybrid quadrupole orbitrap mass spectrometer, equipped with a higher-energy collision induced dissociation cell. Tandem MS was controlled by a data-dependent acquisition. Technical specifics are in the **SI** document.

### Data Repository

The HRMS proteomics data have been deposited to the ProteomeXchange Consortium via the PRIDE^45^ partner repository with the dataset identifier PXD028040.

### Data Analysis

HRMS data were processed in MaxQuant version 1.6.3.3 (Max Planck Institute of Biochemistry) against the mouse proteome (UniProt, downloaded on September 6, 2018). Identified peptides and proteins were filtered to <1% false discovery rate (FDR), calculated against a reversed-sequence decoy database. Common contaminant proteins were annotated and removed from identifications reported in this study. Network prediction was conducted in STRING version 11.5^46^ using gene ontology terms with FDR calculations using the Benjamini–Hochberg procedure. Additional details are in the **SI** document.

## RESULTS AND DISCUSSION

The goal of this study was to integrate neuronal identification with single-cell MS proteomics. Our recent work adapted *in situ*^27,34^ and *in vivo*^33^ capillary microsampling to microanalytical CE-ESI-MS platforms to enable direct proteomics and metabolomics in large, identified cells (~500-to-250 μm diameter) in developing chordate embryos. These instruments preserved cell and embryonic viability, allowing for evaluations of anatomy and whole-organismal behavior.^33^ In 2019, we presented that microanalytical CE-ESI-MS platform was scalable to smaller cells and other functional measurements, including electrical recordings of neuronal activity.^39–41^ This report captures the development, testing, and validation of this platform following the systematic approach outlined in **Figure 1**.

**Figures 1.**
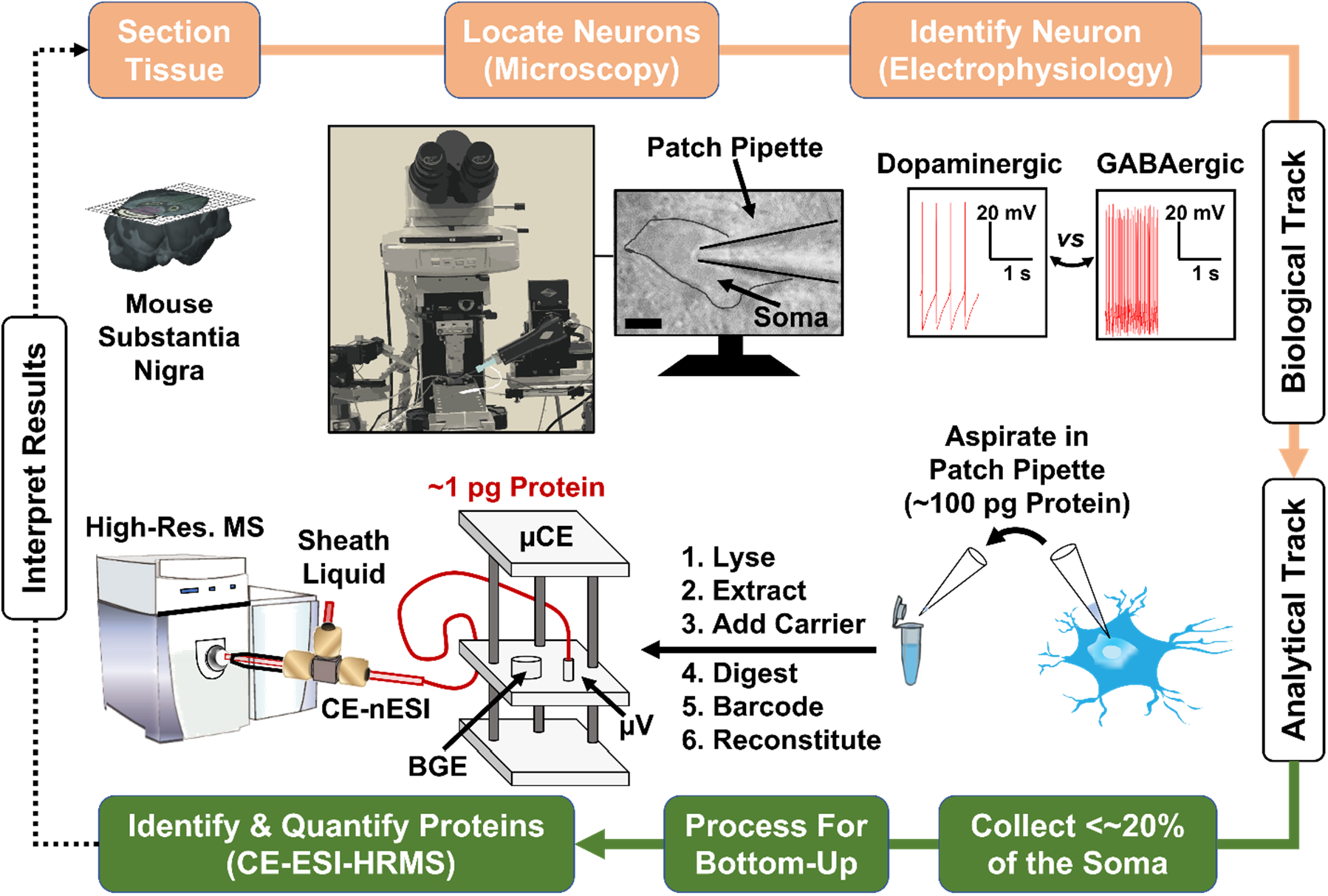
Microanalytical workflow integrating whole-cell patch-clamp electrophysiology and subcellular HRMS proteomics. Dopaminergic (DA) neurons were located based on anatomy and morphology information in high confidence, as confirmed based on their slow pacemaker firing pattern in whole-cell patch-clamp recordings. Example shows fast firing from a nearby GABAergic neuron. See **Figure S1** for higher-resolution data. An ~20–50% volume of the neuronal soma was aspirated into the patch pipette and the collected ~100 pg protein processed for bottom-up proteomics. Ca. ~1 pg of protein, or ~0.25% of the total cell proteome, was analyzed on a custom-built capillary electrophoresis (CE) electrospray ionization (ESI) platform using a high-resolution mass spectrometer (Orbitrap). Key: BGE, background electrolyte vial; μV, sample-loading microvial.

### Technology Development. *Patch-Clamp Electrophysiology*

We began by assessing technical compatibility. We envisioned integration of electrophysiology and CE-ESI-MS to embody a one- or two-step process with respect to the microprobe used for electrical recording and sample collection. **Figure 1** integrates the biological and analytical tracks. For demonstrating proof-of-principle in this study, we selected dopaminergic (DA) neurons as the model, because these cells are readily identifiable based on anatomical location in the brain tissue, morphology and size (~35 μm in diameter), and electrical activity.^2,47,48^ As roughly 70% of neurons in this tissue are dopaminergic,^48^ the substantia nigra pars compacta of the mouse was ideal for our experiments. Acute slices were prepared from the tissue and cultured in modified artificial cerebral spinal fluid to maintain cellular viability.

The neurons were readily identified using an upright microscope. Following standard protocols, the patch pipette was filled with a potassium gluconate-based internal solution, which closely mimics the internal ionic composition of the cell. With the guide of a micromanipulator for precision translation, single putative DA neurons were patched to a Giga-Ohm seal and cell-attached electrical activity was recorded. Technical details are in the **Methods** and the **SI** document. DA neurons were identified by their slow, pacemaker firing rate (2.88 ± 0.42 Hz).^48^ Neighboring cells that exhibited a more rapid firing rate were considered to be GABAergic cells and were not included in this study. Importantly for our study design, microscopy inspection allowed us to identify the DA type with high fidelity after training by patch-clamp electrophysiology. Accurate knowledge of neuron type completed the biological track of our workflow (**Fig. 1**).

To switch to chemical analysis, the next step transformed the patch micropipette to an *in situ/vivo* micro-aspirator. As illustrated in **Figure 1**, we aspirated contents of the neural soma into the patch pipette by applying steady negative pressure from a 3-mL syringe connected to the pipette holder by plastic tubing. Based on microscopic inspection of the shrinking cell, we estimated that ~20–50% volume of the neural soma was collected from each neuron. Our recent experiments using cultured neurons estimated ~500 pg of total protein to be extractable from a single cell.^37^ It follows that approximately 100–250 pg of protein amount was collected from each neuron in this study. Let our calculation conservatively assume collection of ~100 pg or protein from each neuronal soma through the remainder of this report. The microaspirate was ejected into a clean vial, ready for shot-gun proteomics.

Processing of such limited somal protein quantities posed nontrivial challenges. Standard bottom-up proteomics requires extraction and digestion of the proteins, often also calling for chemical reduction/alkylation to deepen the coverage of the detectable proteome. These starting protein amounts, however, were ~1–10 million times smaller than traditionally processed in MS-based bottom-up proteomics, ~1–100 thousand times less than handled from single dissected cells,^29,32,49^ and ~100-times smaller than recently collected from individual cells using capillary microsampling^33,34,50^. As a simplified alternative to automated droplet processing^18–21^, we opted to carry out the processing steps in Lo-Bind vials, common consumables in laboratories (see **Methods**). While this platform ensured simplicity and facilitates methodology adoption in other laboratories, it inherently leads to protein losses on the surfaces of pipette tips and the vial, which we also partially addressed in this study.

A portion of the protein digest was measured using ultrasensitive MS. Our experimental design (**Fig. 1**) leveraged microanalytical CE-ESI-MS for the analyses. Based on our accumulated success, this custom-built platform can measure proteins with high sensitivity, capability for in-column analyte enrichment, scalability to different protein amounts, and quantification using both label-free and multiplexing barcoding strategies. These platforms were reviewed in References^17,23^, and supporting protocols are available in References ^51–53^. The ~100 pg of somal protein digest was reconstituted in 3 μL of 50% ACN containing 0.05% (v/v) AcOH (see **SI** document), which aids sensitivity via field-amplified sample stacking during CE-HRMS.^35^ Following our protocols, ~20 nL or ~0.5% of the sample volume, containing ~1 pg from the somal proteome, was analyzed from aliquots of ~250 nL deposited into a sample-loading microvial (see μV in **Fig. 1**).

Results from this analysis were informative for technical adaptability. CE-HRMS of the somal protein digests returned no protein identification. Closer inspection of the data revealed that the recorded mass spectra were dominated by abundant salt clusters that span across a broad electrophoretic separation window. We ascribe a lack of peptide identifications in these measurements to interferences in electrophoretic separation and electrospray ionization of peptides due to high concentration of nonvolatile salts from the internal solution in the patch pipette. Removal of the nonvolatile salts would support a one-pipette approach (e.g., by C18 binding), as was recently demonstrated in single-neuron analyses using microscale desalting and nanoLC-HRMS.^54^ Using patch recordings, we earlier found neuronal morphology to accurately predict the DA phenotype in our tissue region of interest. Therefore, use of separate patch-pipettes for electrophysiology and neuronal proteomics simplified methods integration in this project.

### Performance Validation

As collection of the protein aspirate required successful patching of the neuron, it was beneficial to explore media compatibility between electrical recording and CE-HRMS. AmBic was selected, being a commonly available, volatile salt, and pH buffer as well as favored sample media also in CE-HRMS proteomics. Experimentally, we found that 50 mM AmBic to closely approximate the osmolarity of the potassium gluconate internal solution (265–285 mOsm). Although electrophysiological recordings from media were of insufficient quality, electrical recordings were sufficient for ensuring patch seal with the neural soma.

For enhanced sensitivity, we refined sample processing and HRMS detection. To facilitate this portion of the study, we prepared standard protein digests from tissues dissected from the substantia nigra pars compacta. The protein digest was suspended in 50 mM AmBic to yield ~100 pg of peptide per ~20 nL fraction, thus each serving a technical replicate of a somal-aspirate equivalent samples. This tissue provided scalable and abundant samples for method development.

Step by step, we advanced CE-HRMS sensitivity, starting with sample processing (**Fig. 2A**). Analysis of ~100 pg of tissue protein digest in 50 mM AmBic returned 20 proteins between technical triplicates, a respectable start. **Table S1A** lists proteins from a replicate. Based on calculated label-free quantitative (LFQ) abundances, which are used as proxy for concentration in CE-MS,^35^ the observed protein abundances spanned 2 decades, a rather limited range (**Fig. 2B**). We recently found elimination of the reduction/alkylation steps to benef sensitivity by minimizing protein losses.^34^ Technical triplicates with this strategy gave 46 proteins, albeit still spanning only 2 log-orders of LFQ range. **Table S1B** tabulates proteins from a replicate. For better sensitivity, we adopted usage of an abundant peptide background to reduce losses for sample proteins.^55^ We chose a tryptic digest of bovine serum albumin (BSA) as the background, as this protein was readily available, affordable, and not anticipated in the brain. After suspending the tissue digest in a media containing BSA digest at 10,000-times higher concentration, technical duplicates of 100 pg of tissue per analysis gave 164 and 225 proteins in, or 263 different proteins cumulatively (see **Tables S1CD)**. Notably, LFQ quantification was expanded over 4 decades, demonstrating the importance of minimizing protein losses during sample preparation.

**Figure 2.**
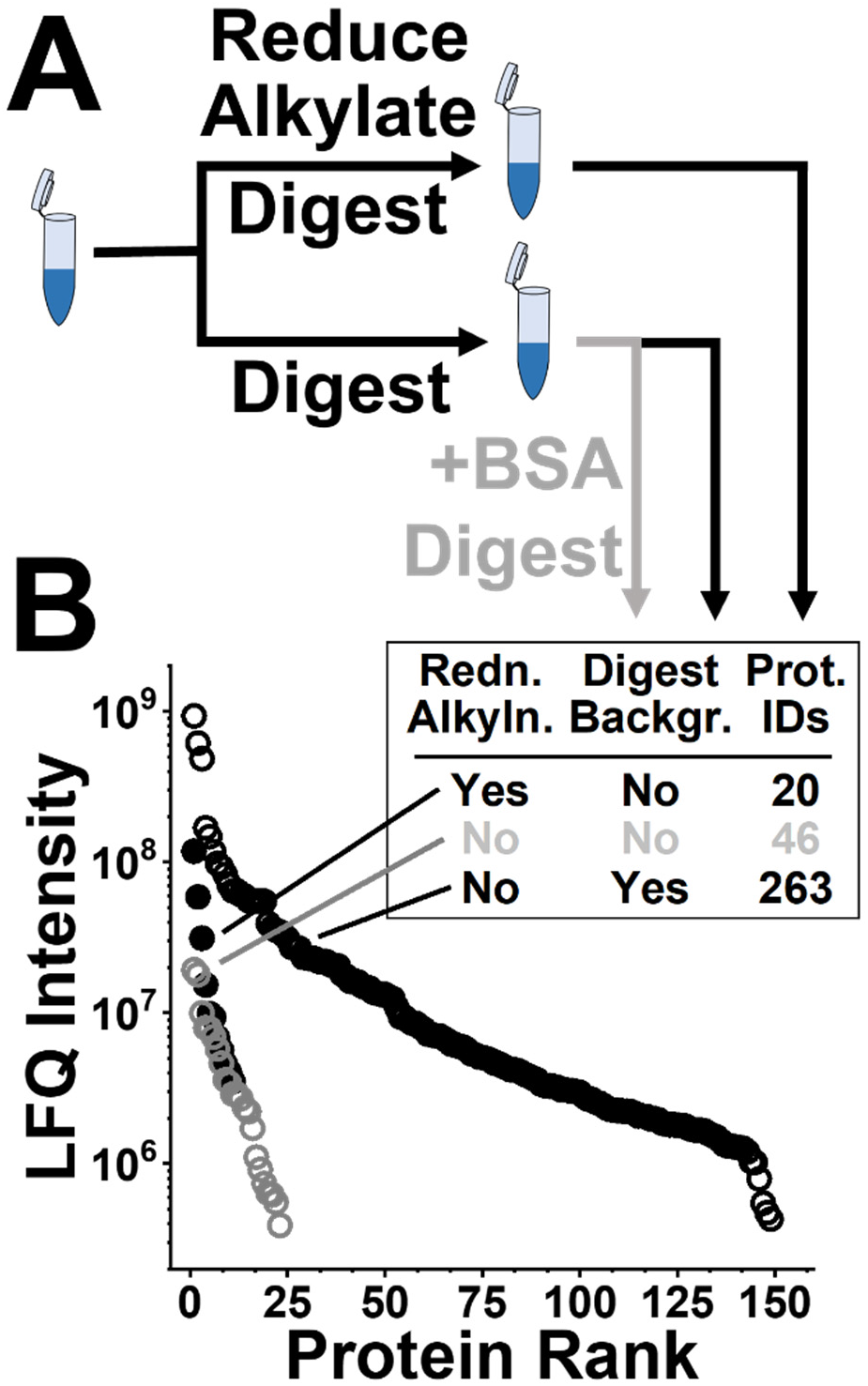
Strategies in sample processing to improve sensitivity. **(A)** Elimination of reduction/alkylation simplified sample processing, while use of abundant peptide backgrounds helped reduce protein losses. **(B)** Comparison of label-free quantification (LFQ) performance between the methods. The carrier protein was digest of bovine serum albumin (BSA), mixed at ~1 ng amount per analysis. As scalable model, ~100 pg of protein digests were measured from the substantia nigra pars compacta, estimating to the somal aspirates.

Mass spectrometric ion detection held the key to added sensitivity (**Fig. 3**). In a typical shot-gun approach, protein identifiction relies on sequencing proteotypic peptides via tandem MS (MS/MS). Multiplexing quantification using designer barcodes, such as tandem mass tags (TMT), recently enabled the incorporation of an abundant carrier channel to enhance the likelihood of ion selection for fragmentation, thus improving detection and quantification sensitivity.^22^ **Figure 3A** shows our strategy for implementation. Tissue protein digests were TMT-tagged and mixed at 1:100 analyte-to-carrier ratio to test the principle.

**Figure 3.**
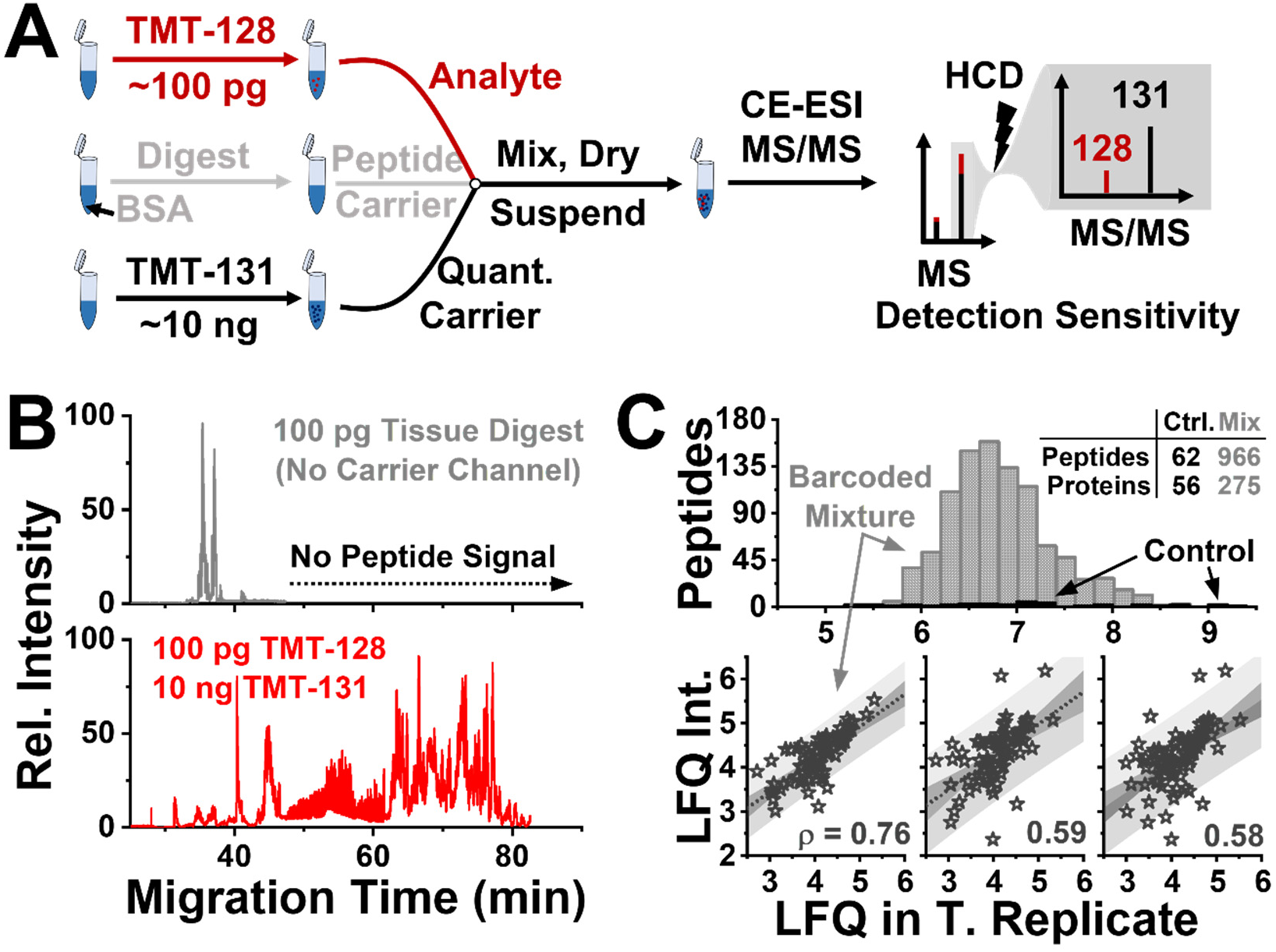
Multiplexing barcoding strategy for enhancing quantitative sensitivity. **(A)** Improvement to ion selection for quantification by supplementing the analyte with a differentially TMT-tagged carrier-proteome (~10 ng of protein digest) at 100-times higher concentration via multiplexing quantification. These proof-of-principle experiments analyzed ~100 pg of TMT-128-tagged and 1 ng of TMT-131-tagged tissue digests from the substantia nigra pars compacta. **(B)** Base-peak electropherograms comparing ion signal abundance from ~100 pg of tissue protein digests before **(top panel)** and after **(bottom panel)** using the TMT-tagged carrier proteome. **(C)** Characterization of label-free quantification for the measured peptides and proteins **(top panel)** and their reproducibility assessment **(bottom panel)** from the control and the barcoded mixture.

Each test analyzed protein amounts equivalent to the somal aspirates. Replicate by CE-MS analyzed ~100 pg of TMT-128-tagged and ~10 ng of TMT-131 tagged protein digest from the tissue. These measurements were compared to 100 pg of (untagged) protein digest as the control, which, too, was suspended in the abundant BSA digest background. **Figure 3B** reveals substantial signal abundance based on the recorded base-peak electropherograms. The detected peptides were assigned to 389 different proteins from triplicates. 275 of these proteins produced quantifiable reporter intensity in the TMT-128 channel, which estimated the somal aspirate. A list of proteins quantified from 3 replicates is provided in **Tables S1EFG**. A total of 91 of these proteins were quantified in all the technical replicates, allowing us to also assess quantitative reproducibility. **Figure 3C** shows linear regression on the measured TMT-128 reporter abundances for the proteins. Moderate to large correlation coefficients (ρ = 0.76, 0.59, and 0.58) revealed appreciable technical repeatability to support quantitative studies.

### Patch-Clamp Proteomics of Single Neuronal Somas

The approach was ready for testing. We identified 3 different DA neurons in the substantia nigra pars compacta and microsampled their somas using the patch pipette. From each neuronal aspirate, proteins were extracted and processed without reduction and alkylation, tagged with TMT-128, mixed with TMT-131-tagged protein digest from the substantial nigra pars compacta, and spiked with a background of BSA digest at ~10,000-times higher concentration (see earlier). For multiplexing carrier, we considered the tissue proteome as a sufficiently close proxy for the DA proteome in these experiments. Future enhancements may benefit from closer alignment between the analyte and carrier proteomes, which is reasonably well obtainable from a small population of pure DA neurons. CE-MS was used to analyze sample amounts containing ~1 pg of somal protein digest in the TMT-128 channel. These protein amounts estimate to ~0.5% of the aspirated protein material, or ~0.25% of the total somal proteome.

These measurements promised rich information on somal proteomes. As our approach by design identified proteins with help from the abundant multiplexing carrier tissue digest (TMT-131 channel), we limited the next portion of data analysis only to proteins that were quantifiable in the analyte channel, viz. had a non-zero ion signal count for the TMT-128 reporter. CE-MS quantified 91, 80, and 95 different proteins in the respective neuronal somas under the tested experimental conditions. The identified proteins are listed in **Tables S1HIJ**. These identifications were aggregated to 157 different proteins between the biological triplicates and are listed in **Table S1K**. Identification and quantification of these proteins did not require functioning probes.

Protein identifications were compared. **Figure 4A** groups the proteins among the somas. These data are provided in **Table S1L.** Abundant cell components were redundantly quantified among the replicates, including various isoforms of actins, heat shock proteins, solute transporters, and tubulins. These molecules include proteins expected to be enriched in neurons, such neurofilament proteins and components of synaptic release machinery. Variable quantification for the other proteins may indicate subcellular biochemical differences within or between the somas. **Figure 4B** evaluates the quantitative variability for 157 different proteins that had quantifiable (non-zero) TMT-128 reporter ion abundance; this channel barcoded proteins from the neuronal soma aspirate. Pearson product moment coefficients of 0.32, 0.54, and 0.61 revealed quantifiable variability between the somal proteome samples. With the CE-MS method tested for moderate-high quantitative reproducibility earlier (recall **Fig. 3C**), we ascribe the observed variability to technical variability between sampling different subcellular components of the soma, innate biological variability between the translational state of the neurons, or their combination. Future studies may leverage our approach to study the molecular underpinnings of the observed quantitative proteomic differences.

**Figure 4.**
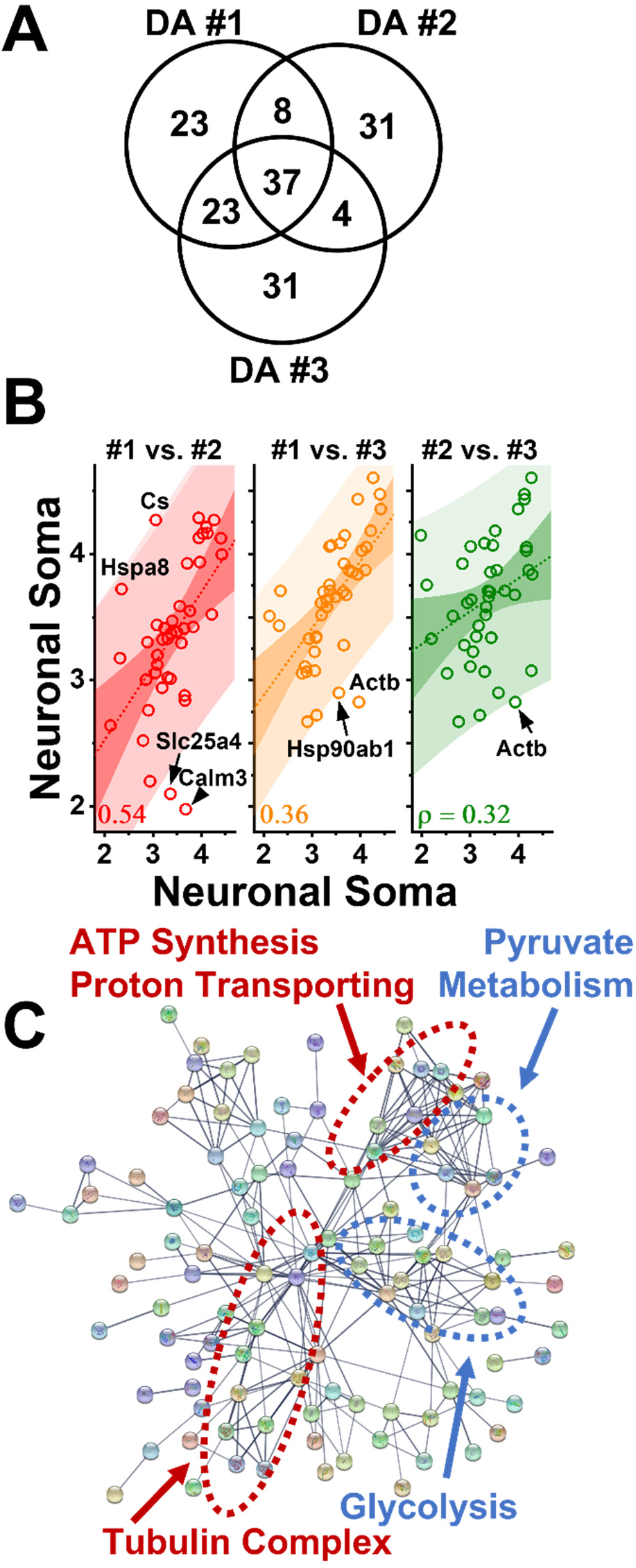
Somal proteomics from single, identified dopaminergic neurons. Ca. 1 ng, or ~0.25% of the somal proteome was analyzed by CE-MS in this work. **(A)** Comparison of 157 different protein quantified between the neuronal somas. **(B)** Correlation analysis of LFQ abundances, revealing detectable proteomic differences between the somal protein aspirates. **(C)** STRING prediction of high-confidence canonical association networks from the quantified proteins. Each dot marks a different protein. Disconnected protein nodes are hidden. A close-up with protein names is shown in **Figure S2**.

The quantitative molecular results could help assess functional differences between the cells. **Table 1** annotates the canonical function of these proteins using STRING analysis. Key biological processes were related to neurofilament assembly and organization and metabolic processes. Molecular processes were enriched in energy production and maintenance. Detection of proteins from the nuclear, mitochondrial, and tubular compartments agreed with subcellular sampling of the soma in this work, which was further supported by detection of proteins with known cytoskeletal localization. **Figure 4B** predicts protein-protein association networks based on canonical gene ontology knowledge of the 157 different proteins that were quantified in either of the 3 biological replicates. Representative associations are marked for energy production, such as adenosinetriphosphase (ATP) synthesis, pyruvate metabolism, and glycolysis. Enrichment of these pathways is in line with known accumulation of mitochondria in neuronal somas. Other known associations, such as formation of tubulin complexes, was also recapitulated in the dataset. A close-up image marking the proteins is provided in **Figure S2**. Combined, these results demonstrated that patch-clamp electrophysiology and CE-nanoESI-HRMS are capable of characterizing protein expression from single DA neuron with high sensitivity to detect neuronal marker proteins.

**Table 1.** STRING functional pathway prediction of proteins quantified among 3 different single dopaminergic neurons. Gene ontology terms were used. Interaction networks were filtered to high confidence (0.700) with no more than 5 interactions shown in the 1^st^ shell. Results are filtered to the top 5 based on enrichment strength calculated between proteins that are observed and expected in the network. Falser discovery Rate (FDR) was calculated using the Benjamini–Hochberg procedure.

| Functional Enrichment | FDR |
| --- | --- |
| **Biological Processes** |  |
| Phosphocreatine biosynthetic process | 0.0494 |
| Neurofilament cytoskeleton organization | 0.0099 |
| Oxaloacetate metabolic process | 0.0118 |
| Mitochondrial ATP synthesis coupled proton transport | 0.0020 |
| ATP biosynthetic process | 0.00017 |
| **Molecular Function** |  |
| L-malate dehydrogenase activity | 0.0415 |
| ATP:ADP antiporter activity | 0.0415 |
| Fructose-bisphosphate aldolase activity | 0.0415 |
| Structural constituent of postsynaptic actin cytoskeleton | 0.0030 |
| Proton-transporting ATP synthase activity | 0.0019 |
| **Cellular Component** |  |
| Tubulin complex | 0.00024 |
| Mitochondrial proton-transporting ATP synthase complex | $1.53 \times 10^{-5}$ |
| Postsynaptic intermediate filament cytoskeleton | 0.0149 |
| Nuclear pore cytoplasmic filaments | 0.0149 |
| Nuclear meiotic cohesin complex | 0.0197 |
| **Subcellular Localization (Compartments)** |  |
| Proton-transporting ATP synthase complex | $9.64 \times 10^{-6}$ |
| Mitochondrial proton-transporting ATP synthase complex | $9.64 \times 10^{-6}$ |
| SLAC complex | 0.0076 |
| Tubulin complexes | $1.06 \times 10^{-9}$ |
| Proton-transporting two-sector ATPase complex | $2.56 \times 10^{-5}$ |

## CONCLUSIONS

In this study, we expanded the analytical toolbox of neuroscience by adapting patch-clamp electrophysiology with subcellular proteomics using bottom-up CE-MS. Since inception in 2019, ^39–41^ the approach has undergone performance improvements that are now sufficient to support neuroscience research. Recording of neuronal activity aided functional characterization of dopaminergic neurons in sections of the substantia nigra pars compacta from the mouse brain. Systematic refinements to sample collection using the patch pipette as well as to processing and detecting the miniscule amounts of proteins allowed us to extend CE-ESI-MS compatibility to conditions of electrophysiology and make nontrivial improvements in detection sensitivity. Analysis of 1 pg of protein digest, or ~0.25% of the total somal proteome quantified ~157 different proteins among the somas of 3 different DA neurons. Correlative analyses of the measured concentrations suggested detectable differences between the proteomic state of these aspirates. The identified and quantified proteins supported prediction of canonical proteinprotein associations with subcellular sensitivity. The approaches reported here mark an exciting milestone development in analytical chemistry and neuroscience. These accumulated results also suggest future sensitivity enhancements possible by automating and downsizing sample processing (e.g., nanoPOTS^18,19^, OAD^20^, and iPAD^21^) and enhancing MS detection sensitivity (e.g., SCoPE^22^ and DDA ladder^38^). Emerging results suggest promising results for single-neuron analyses using microscale desalting and nanoLC-HRMS.^54^ Electrophysiology with subcellular CE-ESI-MS provides previously unavailable information on the functional and proteomic state of neurons, raising a potential to better understand the establishment and maintenance of cell heterogeneity and its role underlying conditions of health and disease.

## ACKNOWLEDGEMENTS

This work was supported by the Arnold and Mabel Beckman Foundation Beckman Young Investigator Grant (to P.N.).

