## Supplementary material for "Patch-Clamp Proteomics of Single Neuronal Somas in Tissue Using Electrophysiology and Subcellular Capillary Electrophoresis Mass Spectrometry": SI Document

Electronic Supplementary Information (SI) for

### EXPERIMENTAL SECTION

**Materials**. All materials were purchased at reagent grade or higher. Dithiothreitol (DTT), ioadoacetamide (IAD), tris-hydrochloric acid (tris-HCl), tris-hydroxy methylaminomethane (Tris-base), potassium chloride (KCl), and sodium hydroxide (NaOH) were obtained from Sigma-Aldrich (St. Louis, MO). Acetic acid (AcOH), formic acid (FA), TCPK-modified trypsin, and ethylenediaminetetraacetic acid (EDTA) were purchased in MS-grade and acetonitrile (ACN), methanol (MeOH), and water were obtained at LC-MS grade (Optima) from Fisher Scientific (Fair Lawn, NJ). Ammonium bicarbonate (AmBic) was from Avantor (Center Valley, PA). Sodium dodecyl sulfate (SDS) was provided by Amresco (Solon, Ohio). Fused silica capillaries (40/90 µm inner/outer diameter) were from Polymicro Technologies (Phoenix, AZ, USA) and used as received. Borosilicate capillaries for electrode capillaries (0.75/1 mm inner/outer diameter) were from Sutter Instrument (Novato, CA). All standards were prepared in 500-µL LoBind protein microtubes from Eppendorf (Hauppage, NY). Tandem mass tag reagents and 1 M triethylammonium bicarbonate (TEAB) were purchased from ThermoFisher Scientific (Waltham, MA).

**Animals and Preparation of Brain Tissue Sections**. All procedures were carried out in accordance with the guidelines of the National Institutes of Health for animal care and use, under the approval of the George Washington University Institutional Animal Care and Use Committee (Approval no. A378). C57Bl6/J mice (Jackson Laboratories) were bred in-house, maintained on a 12-h light/dark cycle, and provided with food and water *ad libitum*. General methods were followed to prepare acute brain tissue sections as described elsewhere.[1] Male mice aged postnatal day (PND) 21–35 were deeply anesthetized with an intraperitoneal injection of ketamine and dexmeditomidine (100 and 0.25 mg/kg, respectively) and perfused with 34 °C NMDG Ringer's solution (in mM): 92 NMDG, 2.5 KCl, 1.2 $NaH_2PO_4$, 30 $NaHCO_3$, 20 HEPES, 25 glucose, 5 sodium ascorbate, 2 thiourea, 3 sodium pyruvate, 10 $MgSO_4$, 0.5 $CaCl_2$.[2] Following perfusion, the brain was rapidly dissected and horizontal slices (220 µm) were prepared in HEPES Ringer's solution (in mM: 86 NaCl, 2.5 KCl, 1.2 $NaH_2PO_4$, 35 $NaHCO_3$, 20 HEPES, 25 glucose, 5 sodium ascorbate, 2 thiourea, 3 sodium pyruvate, 1 $MgSO_4$, and 2 $CaCl_2$)[2] using a vibratome. Slices recovered for 1 h at 34°C in oxygenated HEPES holding solution and then were held in the same solution at room temperature until use.

**Whole-Neuron Electrophysiology and Sample Collection**. Midbrain slices were continuously perfused at 1.5–2 mL ⁄ min with artificial cerebrospinal fluid (aCSF) at 28–32 °C containing (in mM): 126 NaCl, 21.4 $NaHCO_3$, 2.5 KCl, 1.2 $NaH_2PO_4$, 2.4 $CaCl_2$, 1.0 $MgSO_4$, and 11.1 glucose. Patch pipettes for recording were filled with potassium gluconate internal solutions containing (in mM): 117 K-gluconate, 2.8 NaCl, 0.2 MgCl2 5, CaCl2, 20 HEPES, 2 Na-ATP, 0.3 Na-GTP, 0.6 EGTA. Patch pipettes for recording (2–4 MΩ) and protein extraction were backfilled with ~20 µL 50 mM AmBic in water. Dopaminergic neurons were putatively identified based on their location in the lateral portion of the substantia nigra pars compacta and their relatively large size. After obtaining a Giga-Ohm (GΩ) seal, negative pressure was applied to rupture the cell membrane and enter the whole-cell configuration. Steady negative pressure applied at the outlet end of pipette with a syringe to aspirate the contents of the neural soma into the pipette. The neuron was visually inspected during microaspiration under an upright microscope (40× magnification, model FN1, Nikon USA, Melville, NY). Electrophysiological recordings were obtained using a Sutter integrated patch amplifier and Sutterpatch software (Sutter Instruments,

Novato, CA). Upon completion of microaspiration, the pipette was gently removed from the cell and the contents expelled into a 500 μL Eppendorf LoBind vial for bottom-up proteomic analysis. A small portion of substantia nigra tissue containing ~50–100 dopaminergic neurons were also collected for TMT labeling approach for sensitivity enhancement. The samples were frozen on dry ice and stored at –80 °C until analysis.

**Sample Processing for Bottom-up Proteomics**. The collected samples were defrosted, and their protein content was extracted and processed following our established protocols.[3-6] To each subcellular aspirate, we added 5 μL of 50 mM ammonium bicarbonate containing 0.1 μg of trypsin for one-step digestion at 60 °C for 1 h. The resulting neuronal protein digests were vacuum-dried and stored at −80 °C until TMT barcoding.

As a reference for method development and testing, a whole-tissue protein digest was prepared. A portion of the substantia nigra was dissected under the stereomicroscope and lysed in 50 μL of lysis buffer (in mM: 5 EDTA, 20 Tris-HCl, 35 NaCl, and 1% (v/v) SDS), facilitated by periodic ultrasonication for 5 min in an ice-cold water bath. The cell debris was separated by centrifugation at 15,000 × *g* for 10 min at 4 °C, and the supernatant was transferred into a clean 500-μL Eppendorf LoBind vial. Proteins were precipitated in 300 μL of ice-cold acetone at −20 °C overnight. The purified proteins were dried at room temperature, reconstituted in 50 μL of 50 mM ammonium bicarbonate and digested with 1 μg of trypsin for 6 h at 37 °C. The resulting peptides were vacuum-dried and stored at −80 °C until TMT barcoding.

The dried peptide samples were barcoded for HRMS quantification. The reactions were carried out using two channels from a 6plexing kit (TMTsixplex, ThermoFisher Scientific) following vendor protocols. The dried protein digest from each single neuron was reconstituted in 1 μL of 100 mM TEAB and tagged with 1 μL of 85 mM TMT labelling reagent using the TMT-128 channel. The dry peptide sample from the reference tissue was suspended in 20 μL of 100 mM TEAB and tagged with 5 μL of 85 mM TMT label reagent using the TMT-131 channel. Each sample was incubated for 1 h at room temperature for maximal labeling. After incubation, the TMT-128 and TMT-131 reactions were quenched with the addition of 0.5 μL and 2 μL of 5% hydroxylamine, respectively, with incubation for 15 min at room temperature. These tagged samples were mixed such that analysis of each ~20 nL volume of the mixture by CE-MS contained ~100 pg of the TMT-128-tagged digest (analytical channel) and ~10 ng of the TMT-131-tagged digest (carrier channel, used for signal enhancement). For method development, the analytical channel was diluted protein digest prepared from the dissected substantia nigra tissue (see earlier). For single-neuron measurements, the analytical channel contained the protein digest that was prepared from the neuronal soma aspirate. The mixture was vacuum-dried and stored at −80 °C for up to 1 month for analysis.

**Single-cell CE-HRMS.** The dried peptide mixtures were suspended in 3 μL of 50% ACN containing 0.05% (v/v) AcOH, which we previously found efficient for in-column enrichment via field-amplified sample stacking.[5] The labeled peptide samples were measured on the same custom-built micro-loading CE platform that we recently reported.[3] The platform was operated under the same experimental conditions, unless detailed here. The separation CE capillary was coaxially fed into a pulled borosilicate capillary, thus providing a nanoelectrospray (nanoESI) emitter (0.75/1 mm inner/outer diameter) with ~10–15 μm tip aperture. An ~20 nL, containing <1 pg, of protein digest was hydrodynamically loaded into the CE separation capillary and electrophoresed in 25% ACN with 1 M FA (background electrolyte, BGE) in a 100-cm fused silica capillary at ~220 V/cm field strength. Peptides were ionized in the nanoESI interface, which provided a low-flow sheath solution (10% MeOH in 0.05% (v/v) AcOH) throughout the borosilicate capillary emitter, pumped electrokinetically at +1,700 V. The emitter was positioned ~500 µm in front of a mass spectrometer for detection. The CE-nanoESI source was operated in the cone-jet regime for efficient ion generation.[7]

Peptide ions were mass detected, identified, and quantified by HRMS executing data-dependent acquisition for tandem MS. All mass spectrometry measurements in this work were performed on a hybrid quadrupole orbitrap mass spectrometer (Q Exactive Plus, Thermo Scientific) equipped with a higher-energy collision dissociation (HCD) cell for fragmentation. Ions were surveyed between *m/z* 400–1,700 at 35,000 FWHM resolution ($MS^1$) to trigger tandem HRMS on ion signals with peptide-like isotope distribution pattern. DDA parameters were as follows: chromatography peak width (FWHM), 13.0 s; AGC target, $1 \times 10^6$ counts; maximum IT, 50 ms; dynamic exclusion mass tolerance, 5.0 ppm; peptide match, on; exclude isotopes, on; dynamic exclusion, 9.0 s; ion signals excluded below +2 charge state; ion signal intensity threshold, $1.5 \times 10^6$ counts; apex trigger, off. Ions that passed these threshold criteria were selected for HCD with the following settings: maximum IT, 60 ms; isolation window, 1.5 Da; normalized collision energy, 36%; $MS^2$ resolution, 17,500 FWHM; TopN, 20; loop count, 20; fixed first mass, 110.0 m/z; minimum AGC target, $9.2 \times 10^2$ counts. Fragmented ions were dynamically excluded with 5.0 ppm accuracy for 9.0 s before re-consideration for fragmentation.

**Data Analysis**. Primary HRMS–MS/MS data were analyzed in MaxQuant version 1.6.3.3 (Max Planck Institute of Biochemistry) executing the Andromeda search engine[8] against the mouse (*Mus musculus*) proteome (downloaded from UniProt on February 2nd, 2018) as database with the following search parameters: trypsin digestion, up to 2 missed cleavages; variable modification, methionine oxidation; fixed modification, none; precursor mass tolerance ($MS^1$), 20 ppm; fragment mass tolerance ($MS^2$), 4.5 ppm; minimum peptide length, 5. Peptides were filtered to <1% false discovery rate (FDR), calculated against a reversed-sequence decoy database. The reported proteins were grouped based on the closest parsimony principle. Common contaminants were manually removed from the list of proteins that are reported as identified or quantified in this work.

Network prediction was conducted in STRING version 11.5.[9] Gene ontology terms were used. Interaction networks were filtered to high confidence (0.700) with no more than 5 interactions shown in the 1st shell. Results were filtered to the top 5 based on enrichment strength calculated between proteins that are observed and expected in the network. Falser discovery Rate (FDR) was calculated using the Benjamini–Hochberg procedure.

**Data Repository.** The MS proteomics data have been deposited to the ProteomeXchange Consortium via the PRIDE[10] partner repository with the dataset identifier PXD028040.

**Safety Considerations**. Fused silica capillaries and borosilicate capillary emitters, posing potential needle-stick hazard, were handled with attention. Common safety protocols were practiced during the handling of chemicals. All electrically conductive parts of the CE-nanoESI interface were shielded (grounded or isolated) to prevent electrical shock hazard.

### SUPPLEMENTARY FIGURES

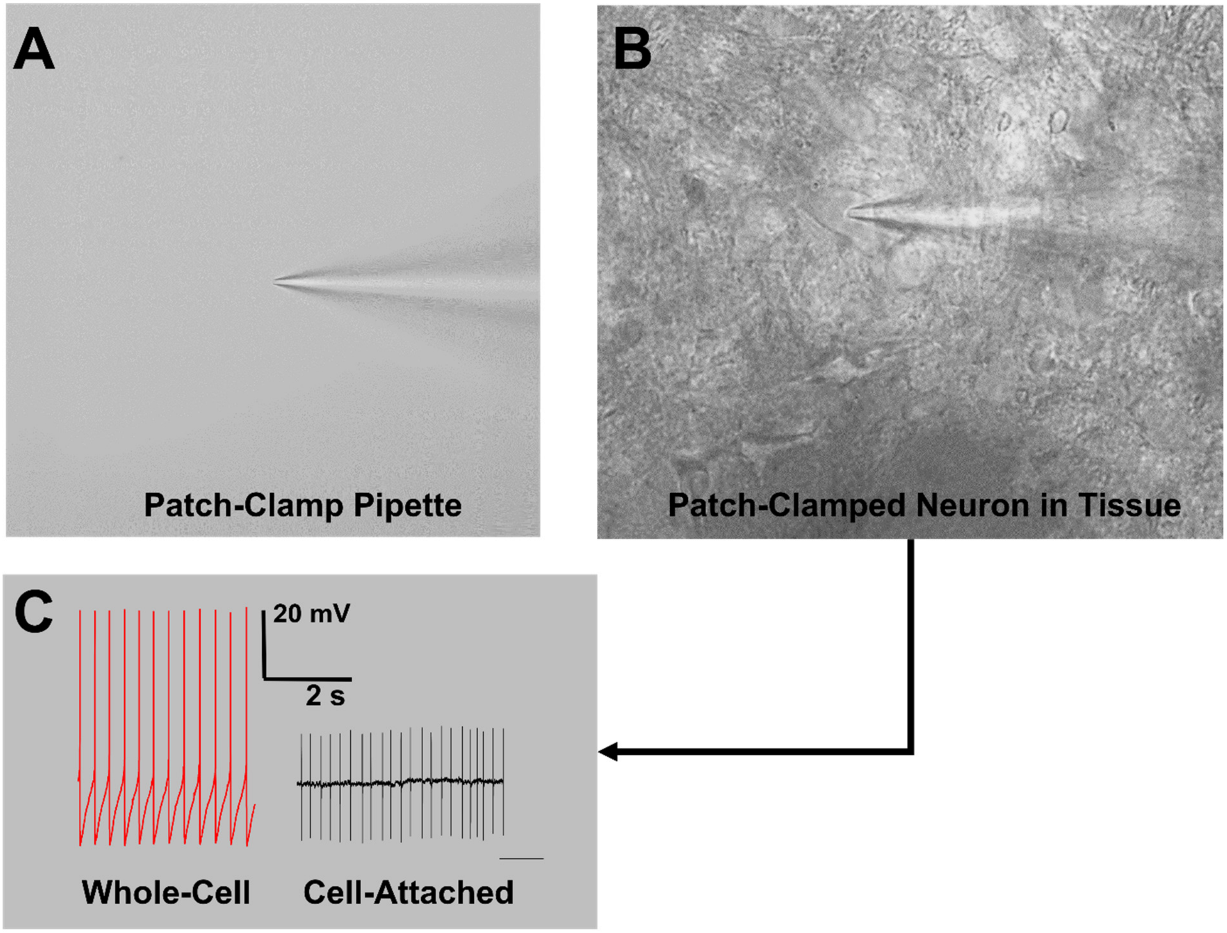

**Figure S1.** Electrophysiological experiments. **(A)** Close-up image of a patch-clamp pipette. **(B)** Example of a patch-clamp pipette contacting a dopaminergic neuron in the substantia nigra. **(C)** Representative examples of whole-cell and cell-attached recordings from a dopaminergic neuron.

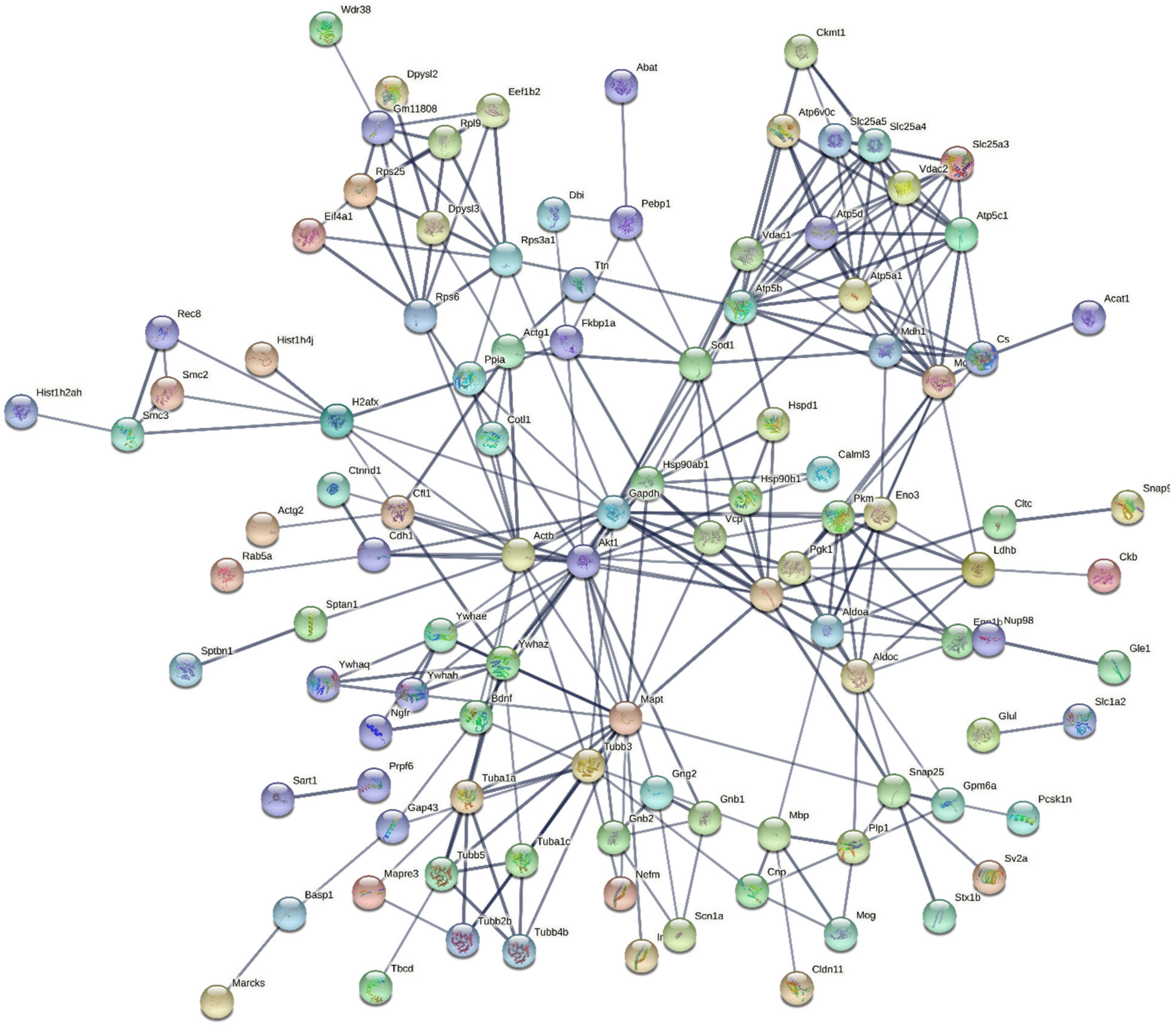

**Figure S2.** STRING functional pathway prediction of proteins that were quantified among 3 different single dopaminergic neurons with non-zero values in the TMT-128 reporter channel measuring that neuronal soma aspirate.
